# Distance to trait optimum is a crucial factor determining the genomic signature of polygenic adaptation

**DOI:** 10.1101/721340

**Authors:** Eirini Christodoulaki, Neda Barghi, Christian Schlötterer

## Abstract

Polygenic adaptation is frequently associated with small allele frequency changes of many loci. Recent works suggest, that large allele frequency changes can be also expected. Laboratory natural selection (LNS) experiments provide an excellent experimental framework to study the adaptive architecture under controlled laboratory conditions: time series data in replicate populations evolving independently to the same trait optimum can be used to identify selected loci. Nevertheless, the choice of the new trait optimum in the laboratory is typically an ad hoc decision without consideration of the distance of the starting population to the new optimum. Here, we used forward-simulations to study the selection signatures of polygenic adaptation in populations evolving to different trait optima. Mimicking LNS experiments we analyzed allele frequencies of the selected alleles and population fitness at multiple time points. We demonstrate that the inferred adaptive architecture strongly depends on the choice of the new trait optimum in the laboratory and the significance cut-off used for identification of selected loci. Our results not only have a major impact on the design of future Evolve and Resequence (E&R) studies, but also on the interpretation of current E&R data sets.

## Introduction

Laboratory natural selection (LNS) has been a popular approach for many years because it is a powerful approach to study adaptation processes under controlled replicated conditions. Polymorphic populations are subjected to different types of stressors (e.g. temperature, desiccation, etc) and monitored for changes in the phenotype of one or several traits that are usually controlled by a large number of loci (i.e polygenic traits). Recently, the analysis of phenotypic change is combined with the analysis of allele frequency changes by contrasting the evolved and ancestral populations (Evolve & Resequence, E&R). The primary goal of such experiments is to uncover the adaptive architecture of the focal trait(s) in populations subjected to certain conditions based on the allele frequency changes during the experiment.

The analysis of different LNS studies revealed contrasting genomic signatures. The most apparent discrepancies between studies are related to the number of putative selection targets and the extent of parallelism across replicates. While some studies detected only a small number of selection targets (Magwire et al. 2012; Mallard et al. 2018), others suggested a polygenic response (Barghi et al., 2019; Jha et al., 2015). While some E&R studies found highly parallel selection response among replicates (Burke et al. 2010; Burke et al. 2016; Graves et al. 2017; Fragata et al. 2018; Mallard et al. 2018; Rebolleda-Gómez and Travisano 2018; Michalak et al. 2019) in other studies selection signatures were much less concordant (Cohan and Hoffmann 1986; Teotónio et al. 2004; Simões et al. 2008; Griffin et al. 2017; Barghi et al. 2019). A particularly striking difference has been observed for natural *Drosophila simulans* populations exposed to the same hot environment. A Portuguese population showed parallel strong selection response at few genomic regions in 5 replicates (Mallard et al. 2018). On the other hand, a population from Florida uncovered highly heterogeneous polygenic selection response among 10 replicates (Barghi et al. 2019). The reason for these different adaptive architectures remains unclear.

In this study we evaluate a potential source of heterogeneity in the adaptive architecture-different selection regimes. We used forward-time simulations to study the influence of the difference between the phenotype of the founder population and the new trait optimum on the adaptive architecture. We show that the distance to the trait optimum has a major influence on the frequency changes of selected alleles and consequently on the underlying adaptive architecture.

## Materials and Methods

We simulated adaptation of a quantitative trait to new optimum in a population of 300 diploid individuals with random mating. The population phenotype (z) is the sum of the phenotypic value of all contributing alleles computed using 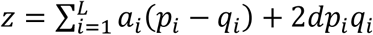 (where α is the effect size, *d* the dominance and *L* the number of the contributing alleles). In all the simulations, no dominance (d=0) and no epistasis were assumed. The phenotypes were mapped to fitness values using a Gaussian function (Fig. S1):

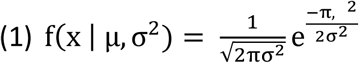

where μ is the mean of the distribution (i.e the new trait optimum) and σ is the standard deviation which is equal to 1 in all of our simulated scenarios. In all simulations, fitness ranged between 0 and 4.5.

To account for linkage, we used 189 *D. simulans* haplotypes for the simulations (Howie et al. 2019). Contributing alleles were randomly distributed across the entire chromosomes 2 and 3 and we used the recombination map of *D. simulans* (Howie et al. 2019). Because *D. simulans* males do not recombine, we divided the recombination rate estimates by two.

We initially simulated a quantitative trait with 100 contributing loci of equal effect size (ES=0.08) and starting frequency (SF=0.2, simulation 1.a in Table 1). Three different trait optima were simulated: close (C), intermediate (I) and distant (D) where the trait optimum was 1, 3, and 5 units away from the phenotype of the founder population. We attempted to mimic an experimental evolution scenario in which a population evolves under three different selection regimes with different intensities (Fig. 1). The values for the three trait optima used in each simulation scenario are shown in Table 1.

**Table 1.** Parameters for the simulations performed in this study. Trait Optimum specifies the new trait optimum values (C: close, I: intermediate and D: distant) that have distance of 1, 3 and 5 units away from the Starting Phenotype. SF: starting frequency, ES: effect size, Nr of loci: Number of contributing loci.

| Simulation | Simulation parameters |  |  |  |  |
| --- | --- | --- | --- | --- | --- |
|  | SF | ES | Nr of loci | Trait Optimum | Starting Phenotype |
| 1.a | 0.2 | 0.08 | 100 | -4 (C), -2 (I), 0 (D) | -5 |
| 1.b | 0.1 | 0.08 | 100 | -5 (C), -3 (I), -1 (D) | -6 |
| 1.c | 0.05 | 0.08 | 100 | -6 (C), -4 (I), -2 (D) | -7 |
| 2 | 0.2 | $\sim \Gamma(0.09, 1)$ | 100 | -2.2 (C), -0.2 (I), 1.8 (D) | -3.2 |
| 3 | $\sim \exp(\lambda = 10)$ | $\sim \Gamma(0.09, 1)$ | 100 | -2.7 (C), -0.7 (I), 1.3 (D) | -3.7 |
| 4 | 0.2 | 0.08 | 50 | -1.4 (C), 0.6 (I), 2.6 (D) | -2.4 |
| 5 | 0.2 | 0.08 | 20 | 0 (C), 2 (I), 4 (D) | -1 |

**Figure 1.**
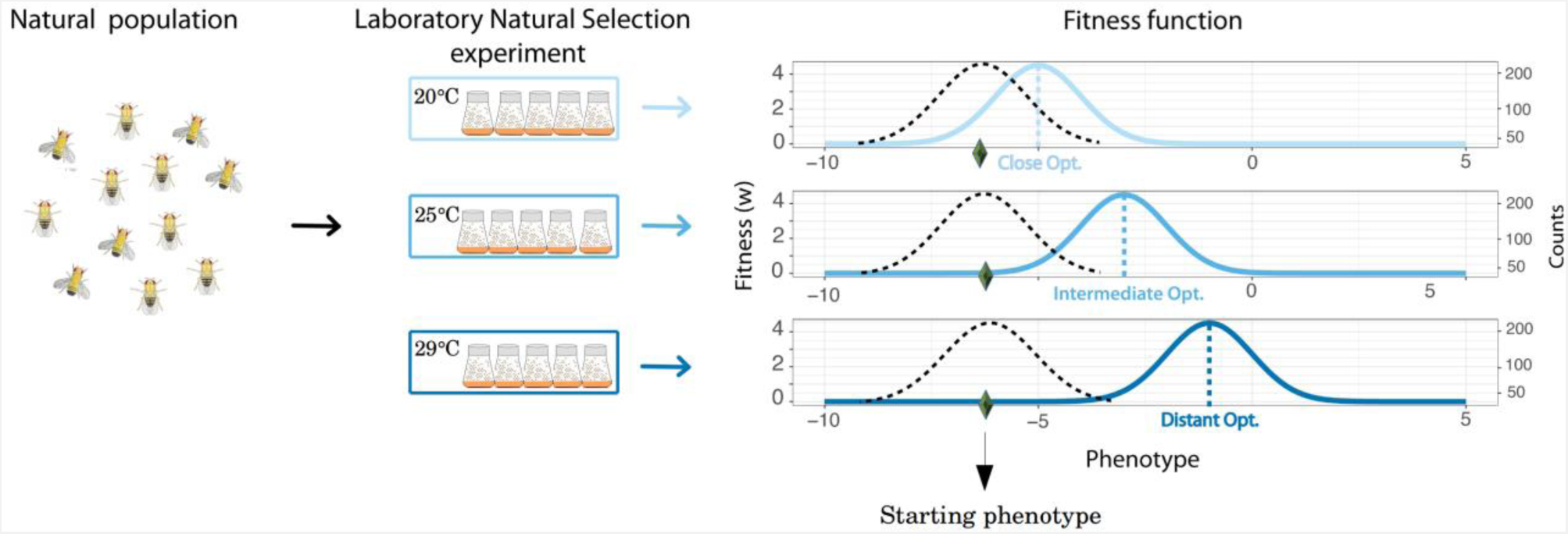
Illustration of the simulated scenarios with varying distances between the phenotype of the founder population and the new trait optimum. We assumed a natural population that adapts to new trait optima that differ in their distance from the phenotype of the founder population. As an example, the corresponding fitness functions are shown. Solid lines show the fitness function and the intensity of shades of blue corresponds to the distance of the new trait optimum: close, intermediate and distant. The black dotted line depicts the distribution of phenotype of the founder population at the start of the experiment and the y-axis on the right shows the number of individuals.

To examine the effect of starting frequency on the inferred adaptive architecture, we simulated a quantitative trait with 100 contributing loci with equal starting frequencies (0.1 and 0.05) and equal effects (0.08, simulations 1.b and 1.c in Table 1).

We also performed simulations matching typical experimental conditions more closely. First, we sampled the effect sizes from a gamma distribution with shape 0.09 and scale 1 (ΓG(0.09,1), simulation 2 in Table 1). The choice of gamma distribution is motivated by findings of QTL mapping studies (Hayes & Goddard, 2001; Hua & Springer, 2018; Mackay, 2010). Second, in addition to sampling the effect sizes from gamma distribution, the starting frequency of alleles was sampled from an exponential distribution with scale 10 (exp(λ=10)), with the majority of alleles starting from low frequencies in the founder population (simulation 3 in Table 1).

We also simulated a quantitative trait with different number of contributing loci (50 and 20) in the founder population (simulations 4 and 5 in Table 1) and equal effect size (0.08).

To enable comparison of simulations with different starting frequencies, effect sizes and number of loci, in each simulation scenario we adjusted the trait optimum such that all populations need to move the same distance from starting phenotype (Table 1) to the new trait optimum, i.e. 1, 3, and 5 units for close, intermediate and distant optima, respectively.

Phenotype and allele frequencies were recorded every 10th generations for 200 generations. For each scenario, we performed 50 replicate simulations using function *qff* in MimicrEE2 (version v194, Vlachos & Kofler, 2018).

To mimic Pool-Seq (Christian Schlötterer et al. 2014), we added a binomial sampling step to the frequencies of the contributing alleles. Selected alleles, i.e. alleles with frequency change more than expected under neutrality, were identified by performing neutral simulations. Neutral simulations were run for 200 generations, identical to the simulations provided in Table 1, but without selection. For each time point we used these neutral simulations to identify selected alleles with frequency changes larger than the 95% cut-off and those with frequency changes less than the 5% cut-off. To infer the adaptive architecture, we classified selected alleles into those with sweep signatures (frequency ≥ 0.9) and small shifts (frequency increase ≤ 10% quantile of allele frequency change distribution at each generation).

## Results

### Distance to trait optimum affects the inferred genetic architecture

We evaluated how the difference between the phenotype of a founder population and the new trait optimum in the experiment influences the inferred adaptive architecture. The genetic architecture is described by the number and allele frequency changes of selected alleles, which deviate from neutral expectations. We classified the allele frequency trajectories as sweep signature and small shifts (Hancock et al. 2010; Berg and Coop 2014; Bourret et al. 2014; Höllinger et al. 2019). In several E&R experiments, the genomic response was studied after about 60 generations of adaptation and strong selection response was identified (Mallard et al. 2018; Barghi et al. 2019; Michalak et al. 2019; Simões et al. 2019). Our standard simulations match this experimental time scale to investigate the adaptive architecture.

We simulated a quantitative trait with 100 linked loci of equal effect (ES=0.08) and starting frequencies (SF= 0.2) (simulations 1.a, Table 1). The computer simulations showed that the distance to the trait optimum affects the inferred adaptive architecture. We observed that the number of selected alleles varies for different trait optima. For example, if the new trait optimum is close (Close scenario in Fig. 2), minor frequency changes are sufficient to reach the new optimum, thus alleles have only small shifts (Table 2). While more pronounced frequency changes occur with increasing distance to the new trait optimum (Fig. 2) that results in more sweep signatures (Table 2).

**Table 2.** Number of selected alleles with sweep signature and small shifts for populations with close (C), intermediate (I) and distant (D) trait optima. SF: starting frequency, ES: effect size, Gamma: gamma distribution, exp: exponential distribution. Simulation parameters are specified in Table 1.

| Simulation | Generation | Sweep Signatures |  |  | Small Shifts |  |  |
| --- | --- | --- | --- | --- | --- | --- | --- |
|  |  | C | I | D | C | I | D |
| 1.a (SF=0.2,ES=0.08,100 loci) | F20 | 0 | 7 | 16 | 19 | 7 | 4 |
| 1.a (SF=0.2,ES=0.08,100 loci) | F40 | 0 | 11 | 21 | 16 | 6 | 3 |
| 1.a (SF=0.2,ES=0.08,100 loci) | F60 | 1 | 12 | 25 | 14 | 6 | 3 |
| 1.b (SF=0.1,ES=0.08,100 loci) | F60 | 1 | 5 | 22 | 18 | 8 | 2 |
| 1.c (SF=0.05,ES=0.08,100 loci) | F60 | 1 | 5 | 19 | 20 | 9 | 2 |
| 2 (SF=0.2,ES ∼ Gamma,100 loci) | F60 | 1 | 1 | 6 | 11 | 11 | 6 |
| 3 (SF ∼ exp,ES ∼ Gamma,100 loci) | F60 | 1 | 2 | 4 | 4 | 5 | 4 |
| 4 (SF=0.2,ES=0.08,50 loci) | F60 | 1 | 9 | 31 | 9 | 2 | 1 |
| 5 (SF=0.2,ES=0.08,20 loci) | F60 | 1 | 11 | 18 | 0 | 0 | 0 |

**Figure 2.**
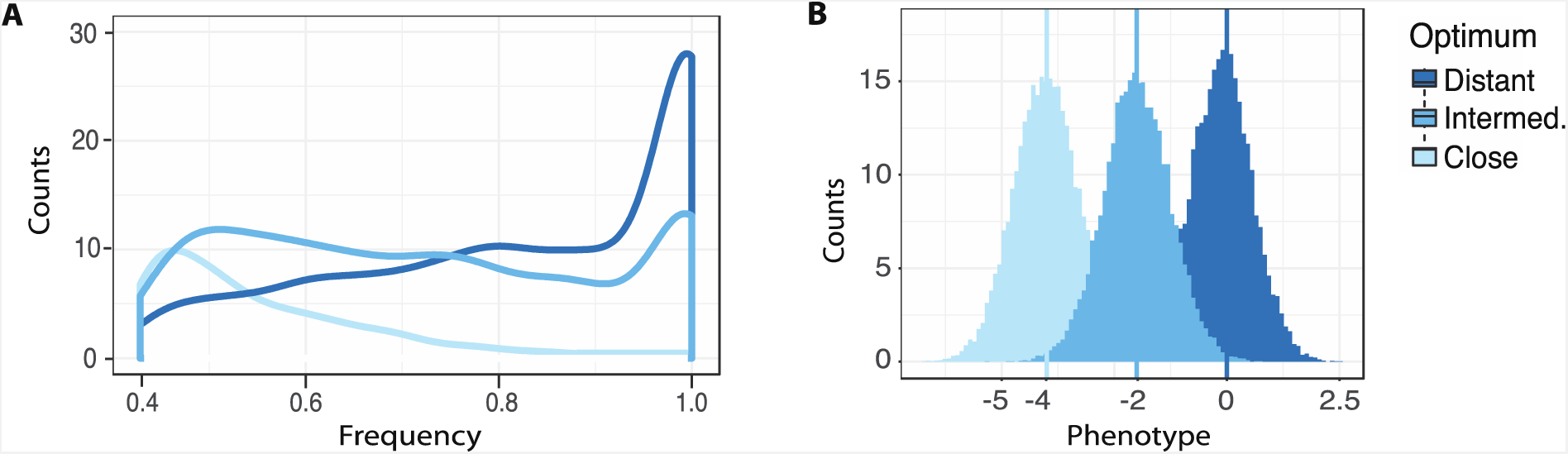
Frequency spectrum of selected alleles (A) and distribution of phenotypes (B) in populations evolved to close, intermediate and distant trait optima after 60 generations. The simulation parameters are specified as 1.a in Table 1.

### Effect of duration of the experiment on the adaptive architecture

The allele frequency trajectories of selected alleles change as the experiment continues for 200 generations (Fig. 3A). The influence of the duration of the experiment is illustrated by the comparison of generations 20, 40 and 60 for populations with different distances to the new trait optimum. Regardless of the duration of the experiment, the distance to trait optimum has a major influence on the genomic signature of adaptation (Fig. 3B). The difference in the number of loci with small shifts and sweep signatures between populations at variable distances to the new trait optimum is persistent at different time points (Table 2).

**Figure 3.**
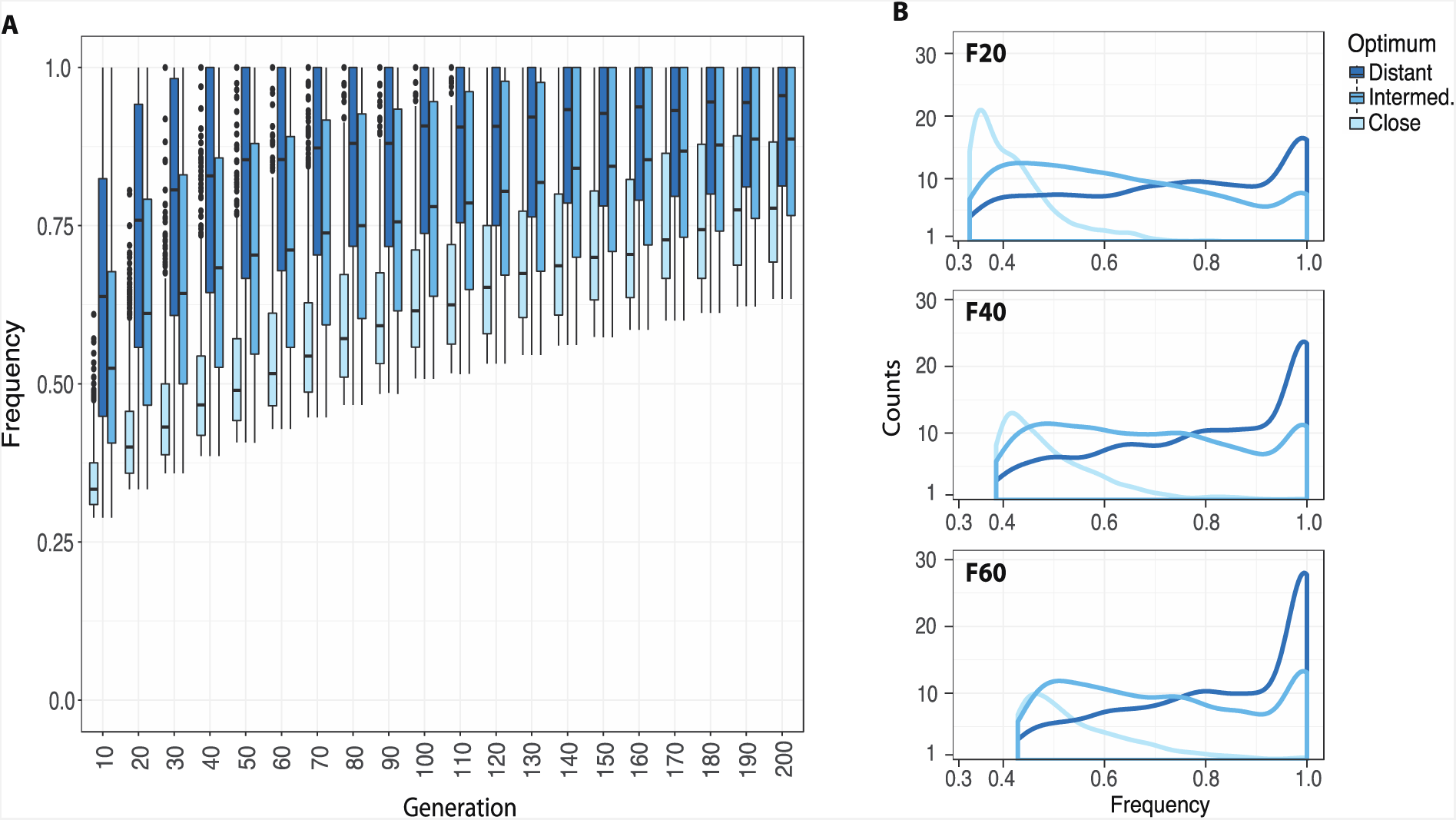
A) Frequency trajectories of selected alleles through 200 generations. B) Frequency spectrum of selected alleles in populations with different distances to the trait optima at generations 20, 40 and 60. The simulation parameters are specified as 1.a in Table 1.

### Effect of allelic starting frequency on the inferred adaptive architecture

We explored the influence of the starting frequency of contributing alleles on the inferred adaptive architecture by simulating 100 loci starting from equal frequencies (0.05, 0.1 and 0.2) and with equal effects (0.08). The starting allele frequencies in the founder population affect the allele frequency changes (Fig. S2) and the evolution of phenotype (Fig. S3). Regardless of the distance to the new trait optimum, populations with higher starting frequencies reach the new trait optimum faster. Higher allele frequencies also result in the loss of fewer alleles due to drift (Fig. S6). This pattern is observed regardless of the distance to the new trait optimum.

Despite the substantial impact of starting frequencies on the allele frequency trajectories and subsequently the inferred adaptive architecture, the influence of the distance to the trait optimum remains fairly similar (Fig. 4). The number of alleles with sweep signatures is higher for populations with more distant trait optima and populations with close trait optimum have more alleles with small shifts (Fig. 4, Table 2). While the number of alleles with sweep signatures and small shifts varies depending on the starting frequencies of alleles in the populations, the distinct patterns of selected loci in populations with different distance to the new trait optimum remain unaffected by the starting frequency (Fig. 4).

**Figure 4.**
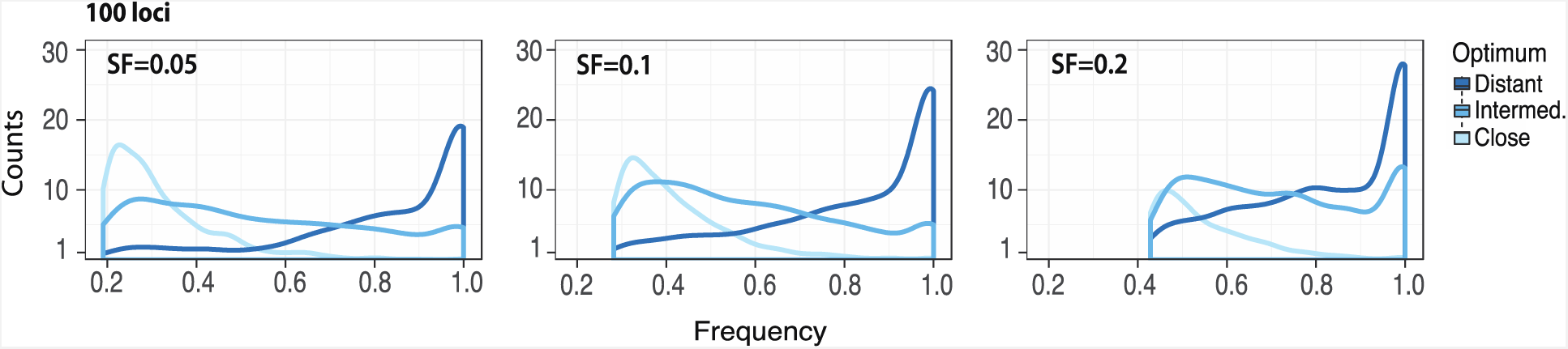
Frequency spectrum of selected alleles with different frequencies (0.05, 0.1, and 0.2) in the founder population and equal effect sizes (ES=0.08) at generation 60. The simulation parameters are specified as 1.a, 1.b and 1.c in Table 1.

### Effect of allelic effect size on the inferred adaptive architecture

We evaluated the influence of allelic effect sizes on the adaptive architecture by simulating a quantitative trait with 100 loci with equal starting frequencies (SF=0.2) and variable effect size. The effect size was sampled from a gamma distribution such that the majority of alleles have small effects and only few are with large effects (simulation 2 in Table 1). Similar to simulations with equal effect sizes across loci, the genomic signatures vary among populations with different distances to trait optimum (Fig. 5). The number of loci with small shifts is quite similar for different distances to the trait optimum (Table 2). However, alleles with sweep signatures, show more prominent differences-distant trait optima result in more loci with sweep signatures (Fig. 5, Table 2).

**Figure 5.**
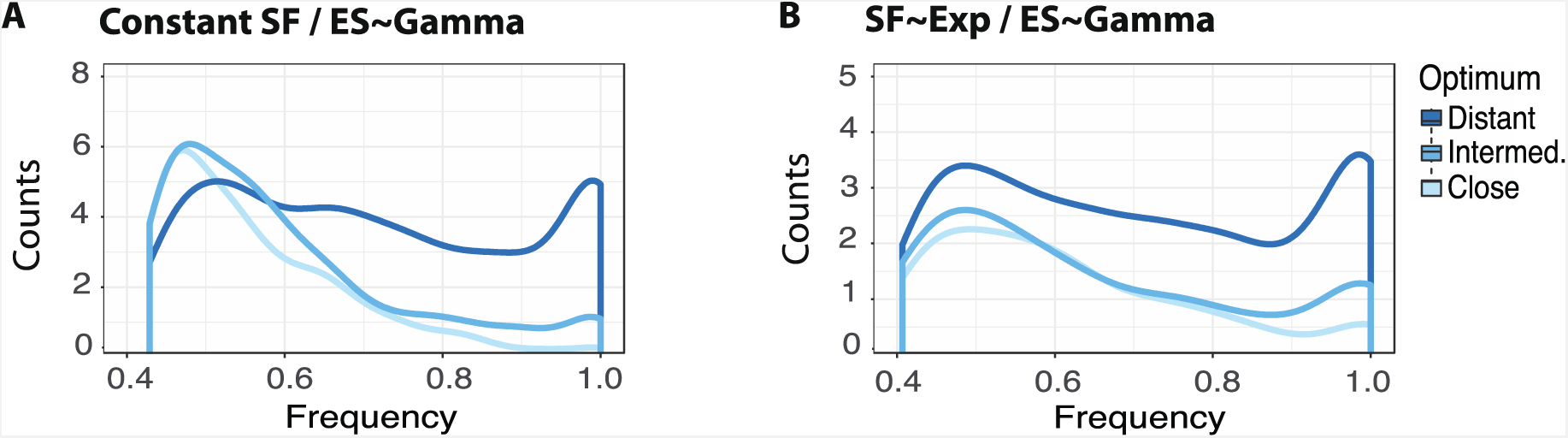
Frequency spectrum of selected alleles with effect sizes sampled from a gamma distribution (Γ(0.09,1)) at generation 60 with A) constant starting frequency (SF=0.2) and B) starting frequencies sampled from an exponential distribution (exp(λ=10)). The simulation parameters are specified as 2 (A) and 3 (B) in Table 1.

Furthermore, we tested the combined influence of variable effect sizes and starting frequencies on the inferred adaptive architecture by simulating effect sizes and starting frequencies sampled from a gamma and exponential distribution, respectively (simulation 3 in Table 1). The exponential distribution mimics a founder population with most selected alleles starting from low frequencies as observed in some experimental evolution studies (Tobler et al. 2014; Barghi et al. 2019). The differences in the inferred genetic responses in populations with different trait optima (Fig. 5B) were not as pronounced as previous simulations. However, as seen in other simulations, sweep signatures were more common for adaptation to an intermediate or distant optimum (Table 2) and thus result in different genomic response among populations.

### Effect of number of contributing loci on the inferred adaptive architecture

We studied the influence of the number of contributing loci on the genomic responses by simulating 100, 50 and 20 loci with equal effect sizes (ES=0.08) and starting frequencies (SF= 0.2) (simulations 1.a, 4 and 5 in Table 1). With more loci in the founder population the new trait optimum is reached faster (Fig. S4). However, the median frequency change of the selected alleles is lower (Fig. S5) because more alleles with smaller frequency changes are required to reach the new trait optimum. This trend is seen regardless of the distance to the new trait optimum.

Our analyses show that the distinct adaptive architectures inferred for different distances to the trait optimum do not depend on the number of contributing loci in the founder population (Fig. 6). Adaptation of populations to distant trait optima is accompanied by higher number of alleles with sweep signatures while alleles with small shifts are observed more when the new trait optimum is closer to the founder population’s phenotype.

**Figure 6.**
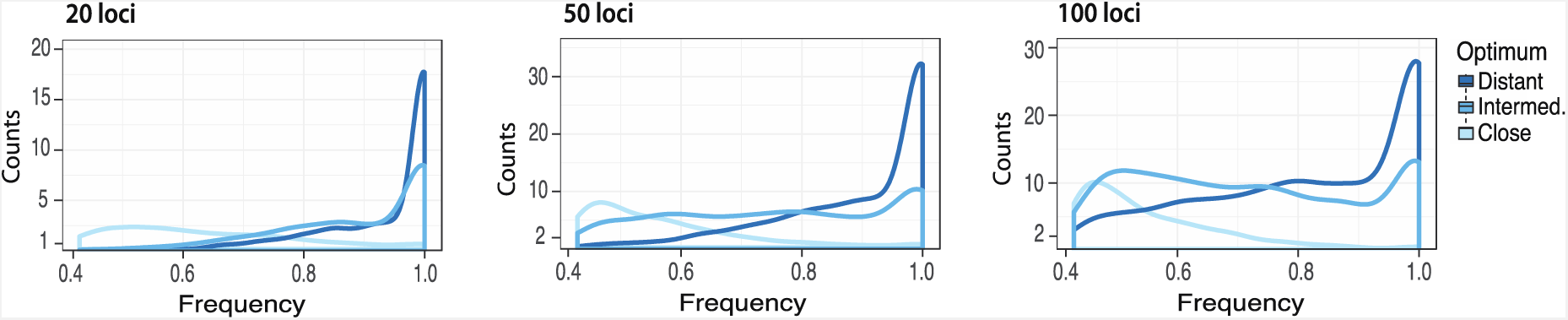
Frequency spectrum of selected alleles at generation 60 with different number of contributing loci (20, 50, and 100) in the founder population with constant starting frequency (SF=0.2) and effect size (ES=0.08). Simulation parameters are specified as 1.a, 4, and 5 in Table 1.

### Effect of neutrality threshold on the inferred adaptive architecture

With the distance to trait optimum affecting the trajectories of allele frequencies and phenotype evolution (Fig. S2, S3) we hypothesized that the power to detect selected loci is also influenced by the distance to the new trait optimum. A more distant trait optimum imposes stronger selection resulting in more prominent allele frequency changes which are more reliably distinguished from neutrally evolving loci.

More stringent criteria for the identification of selected loci will result in the detection of fewer loci. Since the distribution of allele frequency changes differs for the various trait optima, we were interested whether the inferred architecture also depends on the cut-off used to identify selected loci. The choice of neutrality threshold, higher (more conservative) or lower (more liberal) cut-offs, strongly influences the number of identified selected loci, but the distinction between populations adapting to different trait optima remains unaffected (Fig. 7).

**Figure 7.**
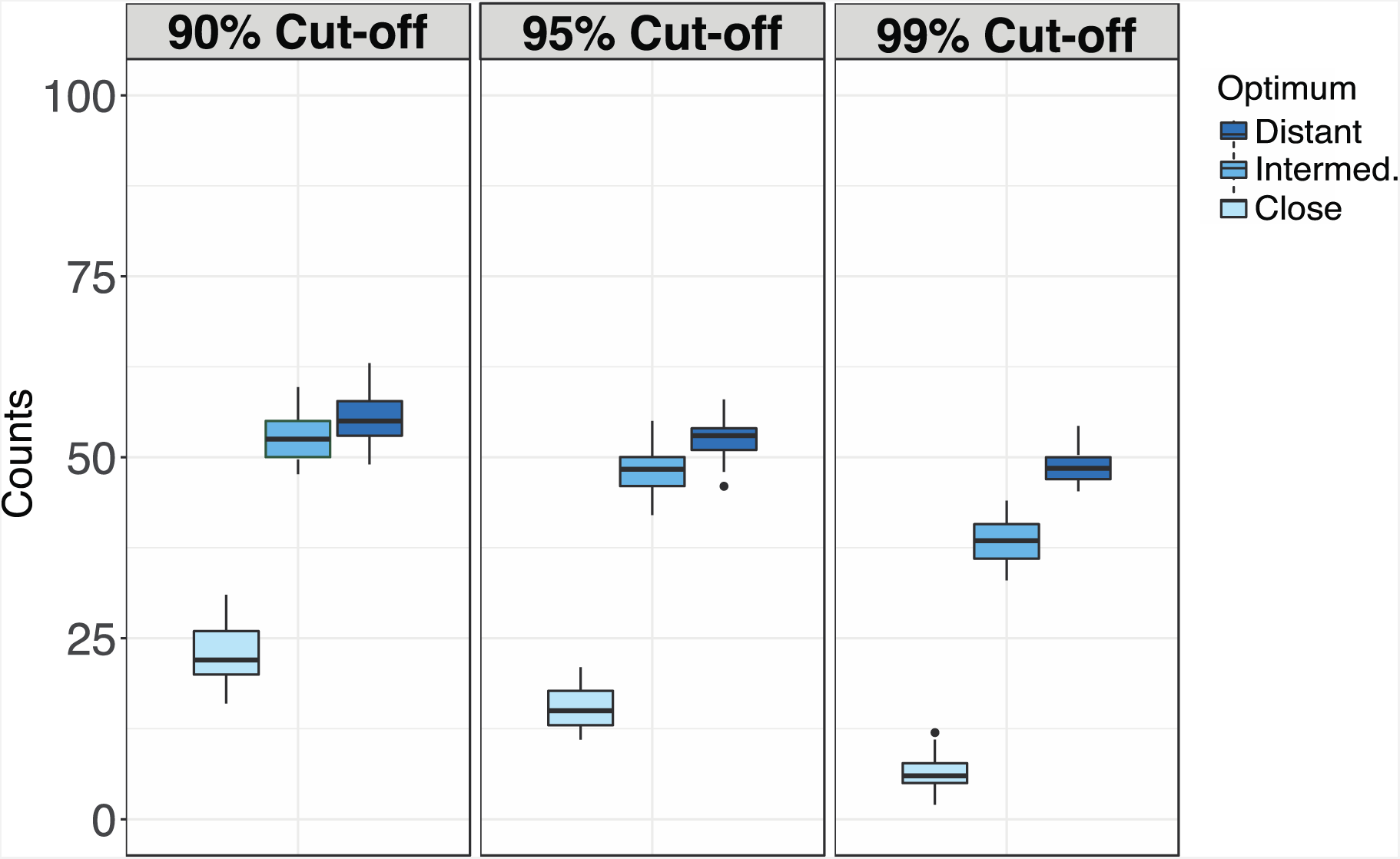
The number of selected alleles at generation 60 with a cut-off of 90%, 95% and 99% determined by neutral simulations. The simulation parameters are specified as 1.a in Table 1.

### Potential mis-classification of selected alleles in experimental populations

Experimental populations are usually maintained at moderate sizes (Matos et al. 2002; Mallard et al. 2018; Barghi et al. 2019; Simões et al. 2019), which results in considerable drift. In particular, low frequency alleles are more prone to loss. In addition to neutral drift, selection also impacts allele frequency changes at neutral and selected alleles. A naive expectation may be that adaptation to close trait optima is based on frequency changes of fewer loci and results in the loss of more beneficial alleles compared to distant trait optima. Simulations with 100 contributing linked loci show, however, the opposite (Fig. 8A). We hypothesized that this pattern results from the interplay of drift and selection, with adaptation to distant optimum causing more intense selection and drift resulting in selected alleles decreasing in frequency. We tested this hypothesis by simulating a quantitative trait with 50 contributing and 50 neutral unlinked loci, and determined the number of selected alleles decreasing in frequency, i.e. alleles with more pronounced frequency decrease than expected under neutrality. These simulations demonstrated that neutral loci in populations with distant trait optimum experience a more pronounced loss than in populations with close optimum (Fig. 8B.2). Consistent with stronger selection in populations with more distant trait optima, their estimated effective population size was considerably smaller than for populations with closer trait optima (Table 3). This result is particularly interesting for experimental studies in which the selected alleles are not known. The natural classification is that the allele with frequency increase is selected. In this case, however, with the decrease in frequency of selected alleles, the alternative allele increases in frequency and would, therefore, incorrectly be classified as the selected allele.

**Table 3.** Effective population size (Ne) computed using neutral alleles at generation 20, 40 and 60 for all trait optima (C: close, I: intermediate, and D: distant), for the simulation with 50 contributing and 50 neutral loci.

| Simulation | Generation | Ne, (50 contributing loci + 50 neutral) |  |  |
| --- | --- | --- | --- | --- |
|  |  | C | I | D |
| SF=0.2, ES=0.08 | F20 | 149 | 73 | 20 |
| SF=0.2, ES=0.08 | F40 | 225 | 126 | 39 |
| SF=0.2, ES=0.08 | F60 | 253 | 150 | 59 |

**Figure 8.**
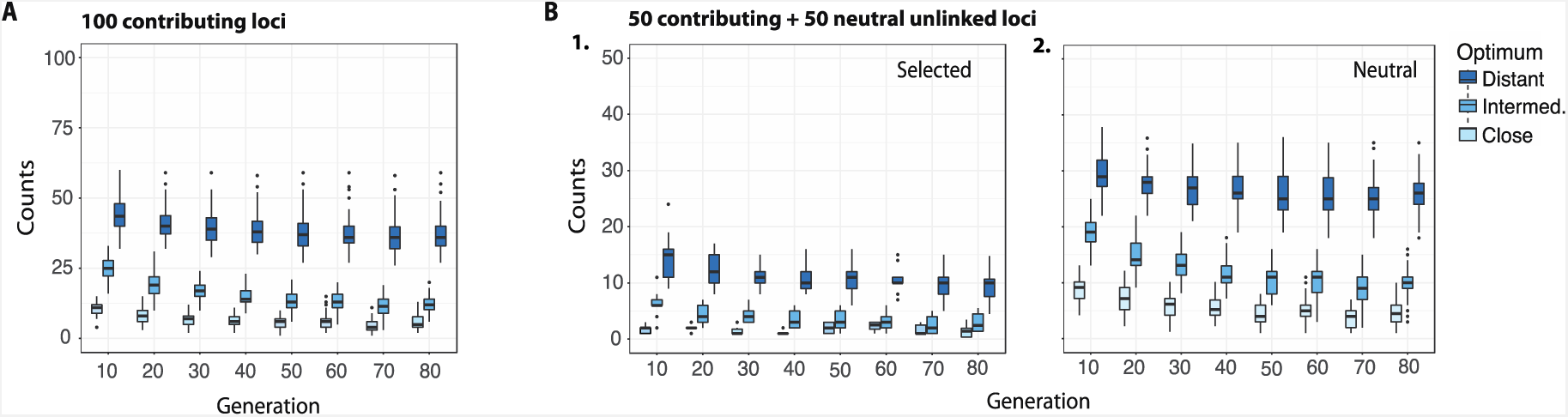
The number of selected alleles decreasing in frequency in simulations with A) 100 linked contributing loci (SF=0.2 and ES=0.08) and B) 50 contributing (SF=0.2 and ES=0.08) and 50 neutral unlinked loci (SF=0.2). The number of selected alleles that decrease in frequency more than expected under neutrality is shown. Decreasing selected alleles are those alleles with frequency changes less than the 5% cut-off from the neutral simulations. The simulation parameters in A are specified as 1.a in Table 1. SF: starting frequency, ES: effect size.

## Discussion

Recently, the combination of experimental evolution with whole genome sequencing (E&R, C Schlötterer, Kofler, Versace, Tobler, & Franssen, 2014; Turner & Miller, 2012; Turner, Stewart, Fields, Rice, & Tarone, 2011) has been a very popular approach to study the adaptive architecture of traits (Teotónio et al. 2004; Burke et al. 2010; Griffin et al. 2017; Mallard et al. 2018; Barghi et al. 2019). While the research goal is well-defined, the impact of the experimental design is not yet fully understood. Previous studies compared the influence of important experimental parameters, such as population size, duration of experiment and number of replicates with computer simulations assuming either adaptation by selective sweeps (Baldwin-Brown et al. 2014; Kofler and Schlötterer 2014) or truncating selection for a quantitative trait (Kessner and Novembre 2015; Lou et al. 2019; Vlachos and Kofler 2019). Very little is known, however, about laboratory natural selection, which aims to mimic natural selection in the laboratory by exposing a polymorphic population to a novel environment. When the allele frequencies of a natural population are preserved in the founder population, LNS does not only identify selection targets, but also provides information about the frequency of the selected alleles in natural populations. Furthermore, LNS can be used to distinguish between polygenic adaptation and the selective sweep paradigm (Barghi and Schlötterer 2019). The choice of the new laboratory environment is frequently an ad hoc decision because of insufficient knowledge about the phenotype and adaptive potential of the founder population. Our study shows that laboratory conditions are not merely a nuisance parameter, but their choice has profound implications.

Consistent with previous studies (Pavlidis et al. 2012; Jain and Stephan 2017; Stetter ID et al. 2018), our simulations demonstrated that several parameters, such as the number of contributing loci, effect size and starting frequency have major influence on the selection response. Independent of these parameters, we show that the distance to the new phenotypic optimum affects the selective response. For more distant trait optima, the selection signature becomes more sweep-like with selected alleles experiencing a substantial allele frequency increase that some reach fixation. For close trait optima, the predominant selection signature is better described by small shifts. Thus, focusing only on the observed allele frequency change, different experiments may result in contrasting conclusions about the underlying adaptive architecture-even when the same founder population was used.

The relevance of the selection regime in the laboratory is nicely illustrated by the evolution of insecticide resistance in the Australian sheep blowfly, *Lucilia cuprina* (McKenzie et al. 1992). Experimental populations that were not mutation limited were exposed to two different selection regimes. In one case the populations were treated with sub-lethal doses of insecticide while in the other one a lethal dose was applied. Sublethal selection regime resulted in the acquisition of many alleles with small effect, like in our simulations of close trait optima. In the other selection regime, the typical major effect loci were identified, matching the loci observed in natural populations.

Our results have more profound implications for the interpretation of selection signatures in LNS experiments than contributing to an improved experimental design. Because the selected phenotype in LNS studies is rarely known, it is impossible to determine the distance of the founder population to the new trait optimum. Consequently, the choice of the laboratory environment will remain an ad hoc decision. The important insight is that, depending on the distance of the trait optimum either sweep-like or small shift signatures can be expected if sufficient genetic variation is present in the founder population. We anticipate that our results also have implications for the interpretation of selection signatures of populations with different origins that evolve in the same laboratory conditions. Populations that are closer to the trait optimum in the laboratory are expected to show more shift signatures, while more distant populations will display sweep-like signatures. We propose that LNS studies which explicitly expose evolving populations to conditions with different trait optima may provide a powerful approach to study the architecture of polygenic adaptation at an unprecedented level.

## Acknowledgements

We thank Nick Barton, Joachim Hermisson, Patrick Phillips and Sam Yeaman for comments and Christos Vlachos for help with MimicrEE2. This work was supported by the European Research Council grant “ArchAdapt” and the Austrian Science Fund (FWF, W1225-B20).

## Data Accessibility

All the information needed to perform the simulations are provided in the Materials and Methods section.

## Author Contributions

Neda Barghi and Christian Schlötterer conceived the study. Eirini Christodoulaki performed the simulations, analyzed and visualized the data. Eirini Christodoulaki, Neda Barghi and Christian Schlötterer wrote the manuscript.

