## Supplemental_Material for "Distance to trait optimum is a crucial factor determining the genomic signature of polygenic adaptation"

### Table of Contents:



| | |
|---|---|
| **Figure S1** | Page 2 |
| **Figure S2** | Page 3 |
| **Figure S3** | Page 4 |
| **Figure S4** | Page 5 |
| **Figure S5** | Page 6 |
| **Figure S6** | Page 7 |

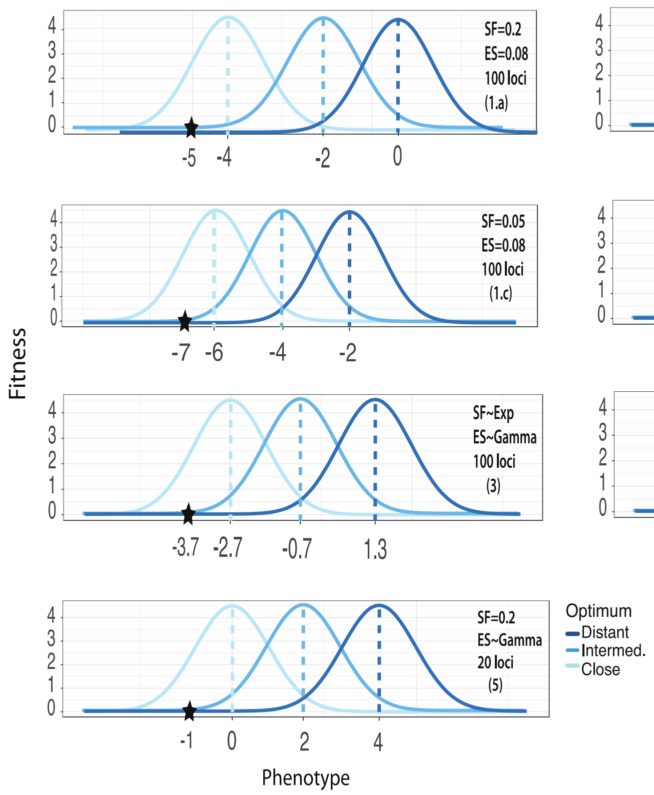


**Figure S1.** Gaussian fitness function to map phenotype to fitness for the different simulation scenarios. Simulation scenarios are specified in parenthesis and the parameters are shown in Table 1. Vertical dashed lines show the new trait optima. The asterisks show the mean phenotype of the founder population (starting phenotype). SF: starting frequency, ES: effect size, Gamma: gamma distribution (Γ(0.09,1)), Exp: exponential distribution (exp(λ=10)).

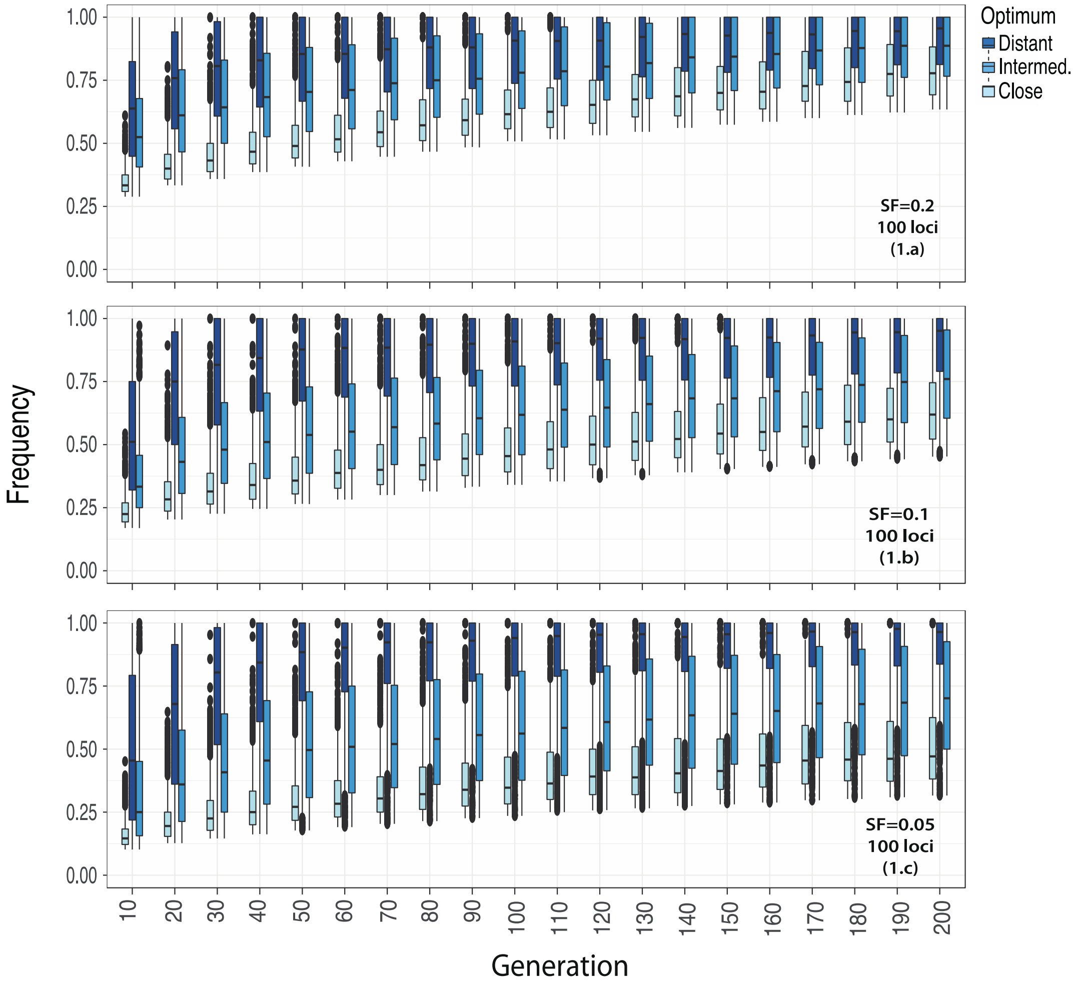


**Figure S2**. Frequency trajectories of selected alleles through 200 generations. A quantitative trait with 100 contributing alleles with equal effect sizes (0.08) and equal starting frequencies (0.05, 0.1, 0.2) was simulated. X-axis shows time in generations while y-axis shows frequency. The simulation parameters are specified as 1.a, 1.b, 1.c in Table 1. SF: starting frequency.

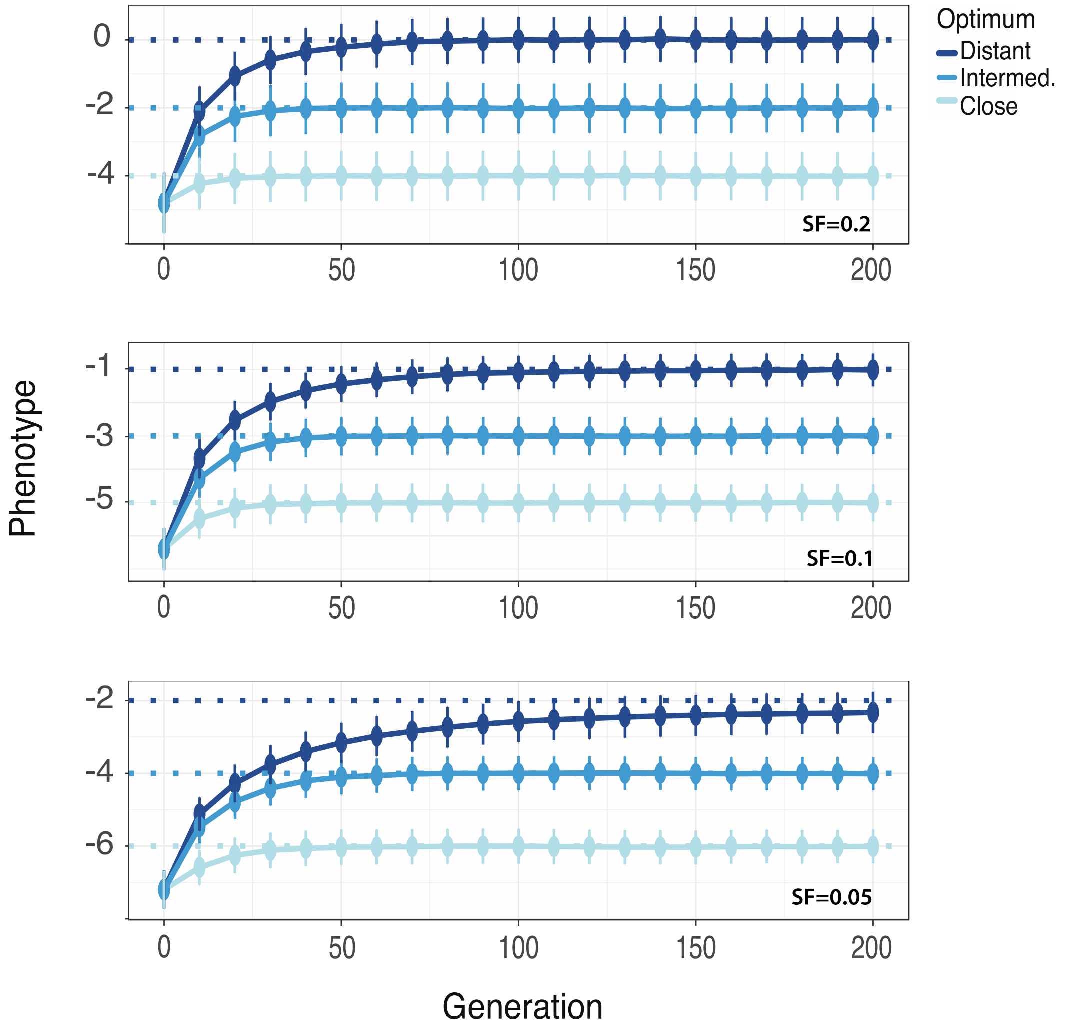


**Figure S3**. Evolution of population phenotype through time in simulations of a trait with 100 contributing loci of equal effect size (ES=0.08) and equal starting frequencies (0.05, 0.1, 0.2). X-axis shows time in generations while y-axis shows phenotype. The three horizontal lines show the three simulated trait optima, the close, the intermediate and the distant. Simulation parameters are specified as 1.a, 1.b and 1.c in Table 1. SF: starting frequency.

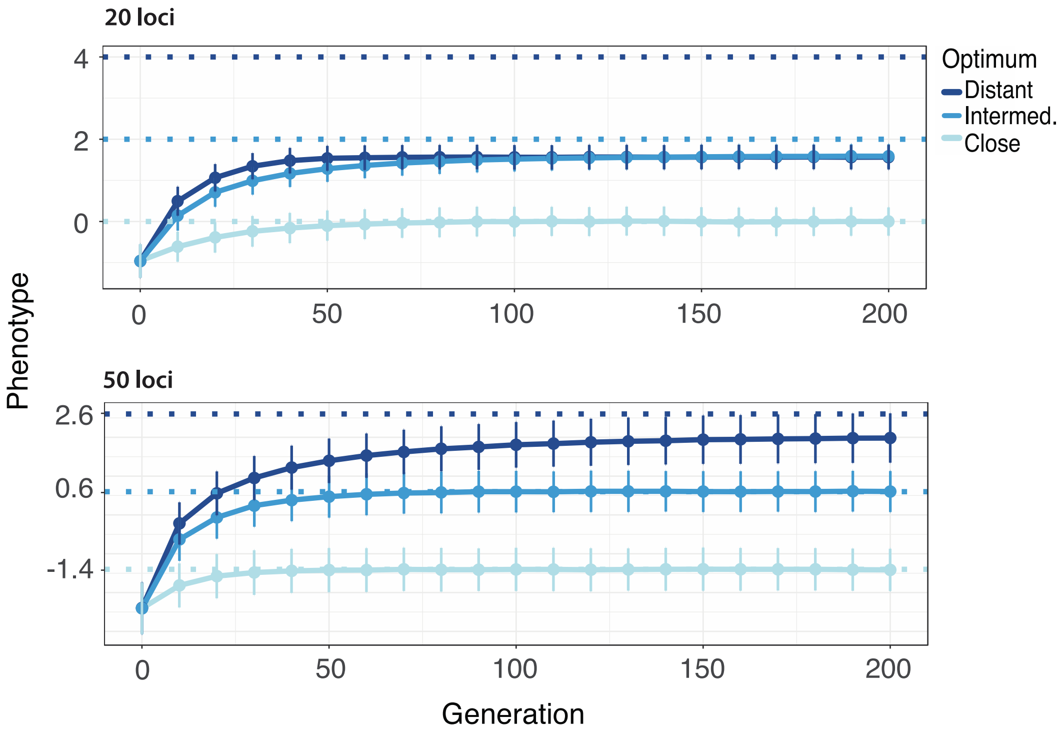


**Figure S4**. Evolution of population phenotype through time in simulations of a trait with 20 contributing loci (first row) or 50 contributing loci (second row) with equal effect sizes (0.08) and equal starting frequencies (SF=0.2). X-axis shows time in generations while y-axis shows phenotype. The three horizontal lines show the three simulated trait optima, the close, the intermediate and the distant. Simulation parameters are specified as 5 and 4 in Table 1.

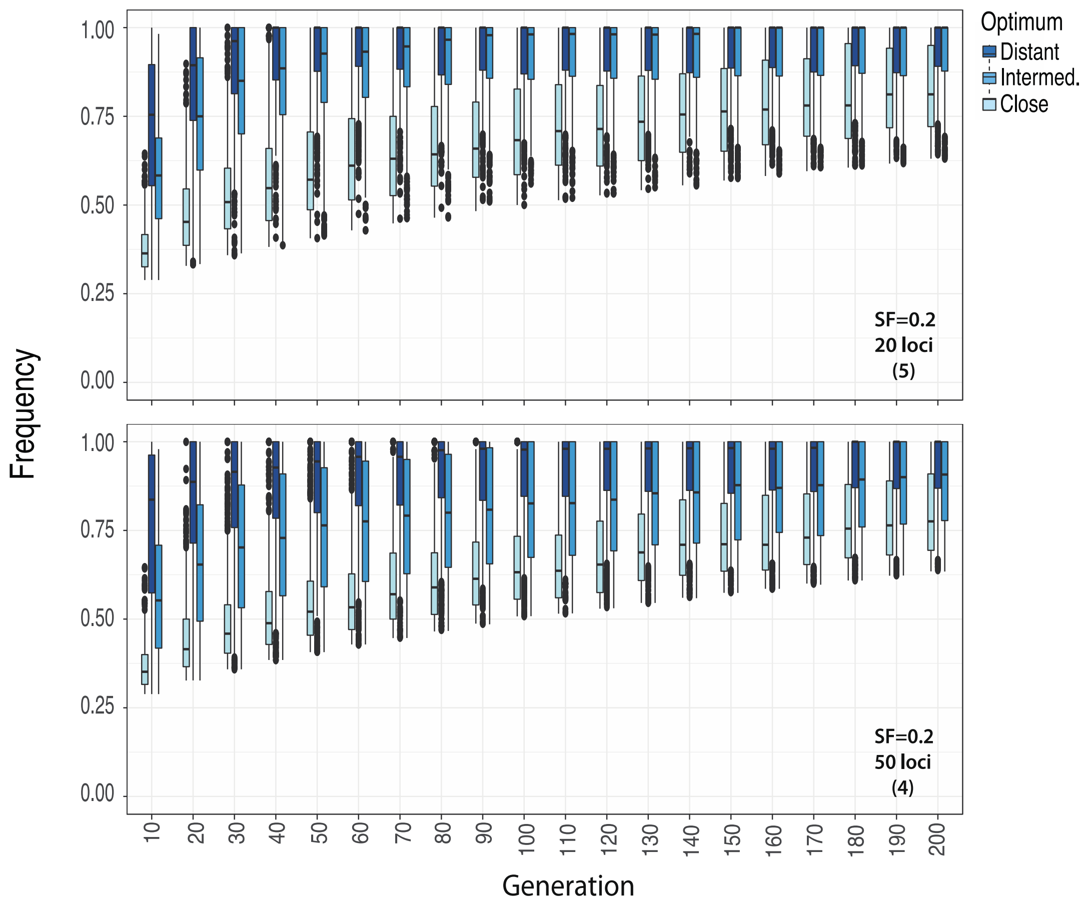


**Figure S5**. Frequency trajectories of selected alleles through 200 generations. A quantitative trait with different numbers of contributing alleles (20 and 50) of equal effect sizes (0.08) and equal starting frequencies (0.2) was simulated. X-axis shows time in generations while y-axis shows frequency. The simulation parameters are specified as 5 and 4 in Table 1. SF: starting frequency.

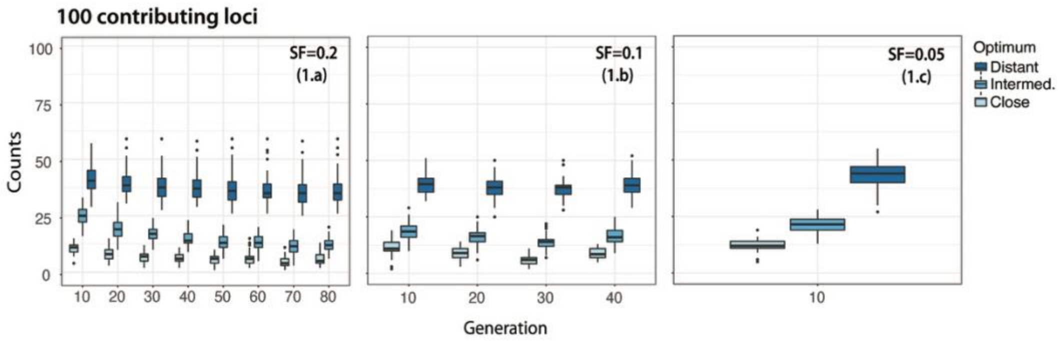


**Figure S6**. Number of selected alleles decreasing in frequency (more than expected by neutrality) in populations with different allelic starting frequencies (0.05, 0.1, 0.2) and different distance to trait optimum. Simulation parameters are shown as 1.a, 1.b, 1.c in Table 1. SF: starting frequency.
